# Identification of functional neural networks of human brains with fMRI

**DOI:** 10.1101/2025.07.07.663492

**Authors:** Jie Huang

## Abstract

The highly evolved human brain comprises numerous functional systems, ranging from essential sensory, motor, attention and memory networks to higher-order cognitive functions like reasoning and language. Although these neural systems and cognitive functions are separately distributed across the entire brain, they are functionally integrated together to perform a task. Decision-making and executive functioning may also be involved in performing the task. While studying task-evoked brain networks is important, investigating whole-brain activity could be crucial for understanding the neural underpinnings of individual behavioral and clinical traits. Even when the brain is not actively engaged in a task, the intrinsic neural activity, i.e., the resting-state (rs) activity, maintains the operations of the brain that involve the acquisition and maintenance of information for interpreting, responding to, and predicting environmental demands. This intrinsic activity is also functionally organized into networks like the brain default mode network. Investigating its whole-brain activity could also be crucial for understanding the neural underpinnings of the brain’s operations at rest. We report a novel data-driven method to objectively and automatically identify functional neural networks (FNNs) across the entire brain for both brain states measured with rs-and task-fMRI, respectively. The identified FNNs characterize the whole-brain activity holistically for each brain state and each individual subject.

---

The non-invasive BOLD-fMRI measures the four-dimensional (3 spatial and 1 temporal) neural activity across the entire brain at large-scale systems level^1-4^. In a typical fMRI study with a spatial resolution of 3×3×3 mm^3^ and a temporal resolution of 2 sec, an activated voxel may contain over one million neurons and its corresponding BOLD signal change measures the activity-induced change from a pooled activity of these million neurons^4^. We conceived the concept of functional area of unitary pooled activity (FAUPA) and developed a novel data-driven method to identify FAUPAs with fMRI^5^. A FAUPA is defined as an area in which the temporal variation of the neural activity is the same across the entire area, i.e., the pooled activity is a unitary dynamic activity. This dynamically unitary activity implies a perfect temporal correlation everywhere within the FAUPA for the activity-induced BOLD response, i.e., the corresponding Pearson correlation coefficient r is equal to 1 for the BOLD responses of any two locations within the FAUPA. FAUPAs have been identified for both rs-and task-fMRI, and their determination was objective and automatic, with no a priori knowledge. The group mean value of r was 0.952±0.004 for the brain state at rest and 0.950±0.002 for the state when performing tasks, showing the dynamically unitary activity within each FAUPA. For each individual subject, using this methodological approach we developed a novel data-driven method to analyze whole-brain activity for each brain state.

## Classification of functional neural networks

We define a FNN as a network in which the temporal variation of the neural activity is the same across the entire network, i.e., all voxels in the network share the same BOLD time signal, resulting in a perfect temporal correlation of this BOLD time signal for all pairwise combinations of these voxels (i.e., the correlation coefficient r=1). The unavoidable physiological and instrumental noises, however, render this correlation unperfect, resulting in r<1. As these noises may vary from time to time and from location to location, their effects may also vary from network to network. For each FNN, its neural activity is reflected in the mean BOLD time signal averaged over all voxels within that FNN. We classify FNN into four categories: (a) FNNs in category 1 (C1), the neural activity in each FNN is very similar across all voxels within that FNN. For each FNN, the minimum r value of the temporal correlation of the mean BOLD time signal with each voxel’s time signal is larger than 0.85 (*p*<3.6×10^−47^ for N=288) for all voxels within that FNN; (b) FNNs in C2, the neural activity is strongly correlated across all voxels within each FNN (i.e., r>0.75 and *p*<4.1×10^−37^); (c) FNNs in C3, the neural activity is highly correlated across all voxels within each FNN (i.e., r>0.65 and *p*<2.7×10^−28^); and (d) FNNs in C4, the neural activity is correlated across all voxels within each FNN (i.e., r>0.55 and *p*<1.0×10^−20^). We developed a novel data-driven method to objectively and automatically identify FNNs successively, starting from the identification of all FNNs in C1, then all FNNs in C2, followed by all FNNs in C3, and finally all FNNs in C4. Table 1 tabulates the identified FNNs in C1 for both brain states for a representative subject, and the supplementary tables 1-3 tabulate the identified FNNs in C2, C3 and C4, respectively. Similar results were obtained for each of the rest eight subjects (data were not presented).

**Table 1.** Functional neural networks in category 1 determined for both brain states for the representative subject. #: number; r_max: maximum r; r_min: minimum r; mn: mean r value averaged over all voxels within each FNN; and sd: the corresponding standard deviation of these r values.

| Task |  |  |  |  |  | resting-state |  |  |  |  |  |
| --- | --- | --- | --- | --- | --- | --- | --- | --- | --- | --- | --- |
|  |  | Pearson Correlation Coefficient r |  |  |  |  |  | Pearson Correlation Coefficient r |  |  |  |
| FNN | # of voxels | r_max | r_min | r_mn | r_sd | FNN | # of voxels | r_max | r_min | r_mn | r_sd |
| 1 | 229 | 0.975 | 0.850 | 0.896 | 0.029 | 1 | 546 | 0.957 | 0.850 | 0.887 | 0.025 |
| 2 | 193 | 0.962 | 0.850 | 0.890 | 0.027 | 2 | 180 | 0.978 | 0.851 | 0.910 | 0.035 |
| 3 | 155 | 0.963 | 0.851 | 0.900 | 0.027 | 3 | 167 | 0.958 | 0.850 | 0.891 | 0.026 |
| 4 | 102 | 0.965 | 0.853 | 0.900 | 0.032 | 4 | 148 | 0.968 | 0.851 | 0.896 | 0.030 |
| 5 | 86 | 0.975 | 0.856 | 0.903 | 0.033 | 5 | 135 | 0.955 | 0.850 | 0.892 | 0.028 |
| 6 | 85 | 0.960 | 0.851 | 0.895 | 0.023 | 6 | 117 | 0.956 | 0.854 | 0.895 | 0.028 |
| 7 | 80 | 0.980 | 0.852 | 0.914 | 0.036 | 7 | 111 | 0.976 | 0.850 | 0.911 | 0.036 |
| 8 | 61 | 0.980 | 0.851 | 0.913 | 0.042 | 8 | 111 | 0.965 | 0.852 | 0.903 | 0.029 |
| 9 | 60 | 0.962 | 0.850 | 0.905 | 0.031 | 9 | 103 | 0.951 | 0.853 | 0.891 | 0.025 |
| 10 | 55 | 0.959 | 0.851 | 0.890 | 0.029 | 10 | 98 | 0.955 | 0.850 | 0.892 | 0.023 |
| 11 | 54 | 0.952 | 0.857 | 0.901 | 0.026 | 11 | 92 | 0.962 | 0.852 | 0.889 | 0.026 |
| 12 | 50 | 0.958 | 0.856 | 0.898 | 0.029 | 12 | 87 | 0.942 | 0.850 | 0.889 | 0.023 |
|  |  |  |  |  |  | 13 | 72 | 0.966 | 0.854 | 0.902 | 0.032 |
|  |  |  |  |  |  | 14 | 69 | 0.970 | 0.850 | 0.905 | 0.033 |
|  |  |  |  |  |  | 15 | 68 | 0.963 | 0.858 | 0.908 | 0.029 |
|  |  |  |  |  |  | 16 | 66 | 0.953 | 0.853 | 0.891 | 0.026 |
|  |  |  |  |  |  | 17 | 64 | 0.963 | 0.857 | 0.895 | 0.026 |
|  |  |  |  |  |  | 18 | 63 | 0.971 | 0.850 | 0.913 | 0.031 |
|  |  |  |  |  |  | 19 | 60 | 0.985 | 0.858 | 0.918 | 0.036 |
|  |  |  |  |  |  | 20 | 60 | 0.943 | 0.852 | 0.896 | 0.025 |
|  |  |  |  |  |  | 21 | 57 | 0.970 | 0.857 | 0.914 | 0.034 |
|  |  |  |  |  |  | 22 | 52 | 0.965 | 0.850 | 0.903 | 0.033 |
| mean | 101 | 0.966 | 0.852 | 0.900 | 0.030 | mean | 115 | 0.962 | 0.852 | 0.900 | 0.029 |
| SD | 59 | 0.009 | 0.003 | 0.007 | 0.005 | SD | 103 | 0.011 | 0.003 | 0.009 | 0.004 |

The size of brain varies from subject to subject. The total number of voxels that cover the entire brain ranges from 22005 to 29249 with mean ± SD = 25301 ± 2229. For each brain state and each subject, we computed the distribution of the correlation coefficient r of voxel’s time signal for pairwise combinations of all voxels. The left nine subplots in Fig. 1 compare the histogram of this r distribution between the two brain states and across the nine subjects. The variation in these histograms demonstrates, to a certain degree, a variable whole-brain activity from state to state and from subject to subject. For each brain state and each subject, both the total number of FNNs in C1 and their corresponding total voxels in these FNNs also varied substantially from state to state and from subject to subject (Fig.1, right). Similar results were obtained for the FNNs in the other three categories (Suppl. Fig. 1). The total number of FNNs in all four categories ranged from 65 to 138 with mean ± SD = 105 ± 24 for the brain state of performing tasks and from 61 to 188 with mean ± SD = 116 ± 42 for the resting state. The corresponding total number of voxels in all FNNs, relative to the total number of voxels of the entire brain, ranged from 60.4% to 90.75% with mean ± SD = 78.43% ± 8.88% for the former state and from 70.6% to 94.15% with mean ± SD = 83.51% ± 7.24% for the latter state. For each FNN, there are four parameters that characterize the FNN: the minimum r value among all voxels, the maximum r value, the mean r value averaged over all voxels, and its corresponding SD. For each brain state and each subject, we computed the mean values of these four parameters averaged over all FNNs for each category. For each FNN category and each parameter, this mean value showed a remarkable similarity between the two brain states and across the nine subjects (Fig. 2).

**Fig. 1.**
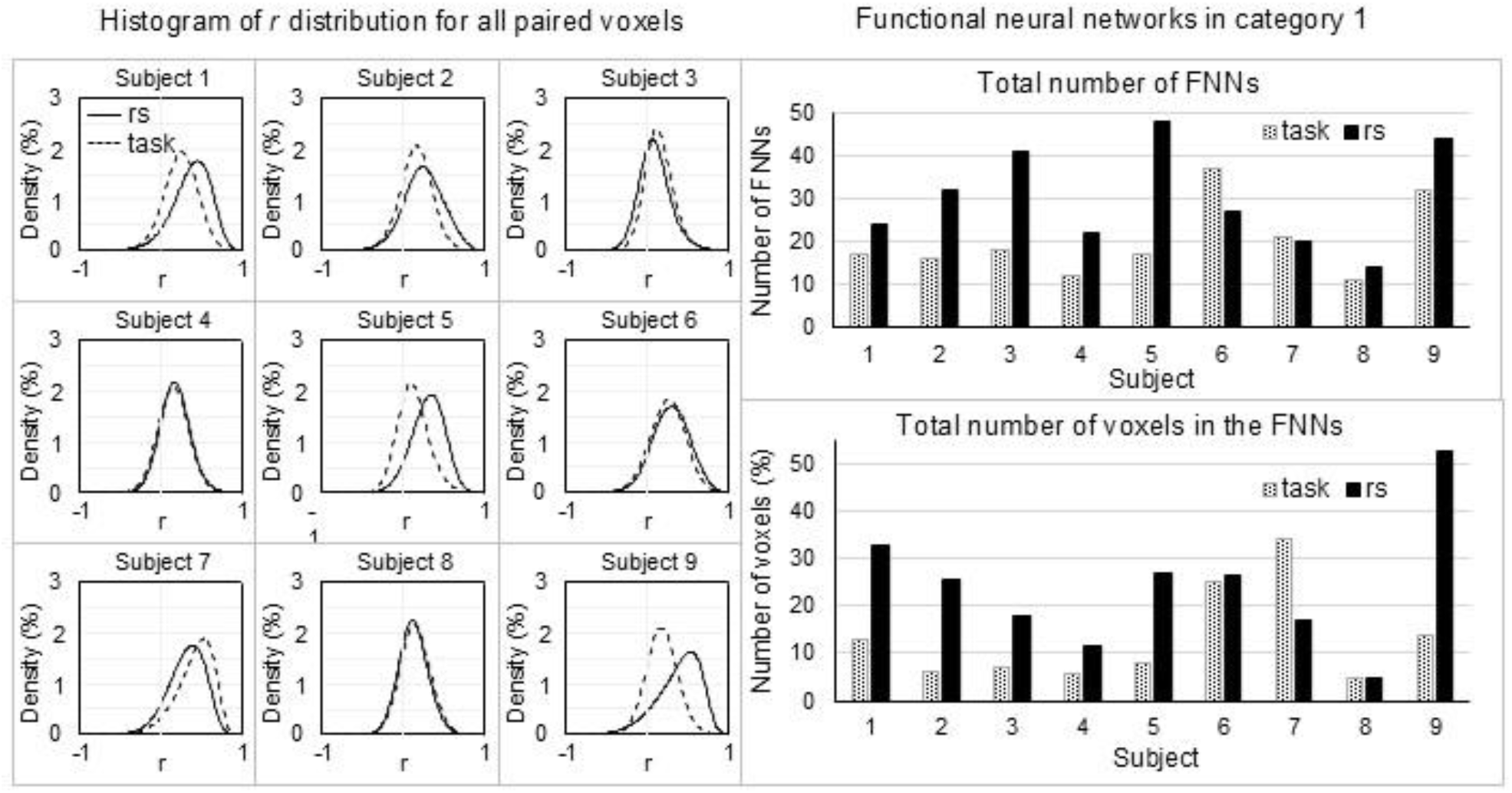
Comparisons of the whole-brain activity and of the identified FNNs in category 1 between the two brain states and across the nine subjects. The left nine subplots illustrate the histogram of r distribution that showed a substantial variation between the two brain states for five subjects. The right top plot illustrates the total number of FNNs in C1 that varied substantially between the two brain states and across the nine subjects. The right bottom plot illustrates their corresponding percentage of total number of voxels in these FNNs relative to the total number of voxels of the entire brain.

**Fig. 2.**
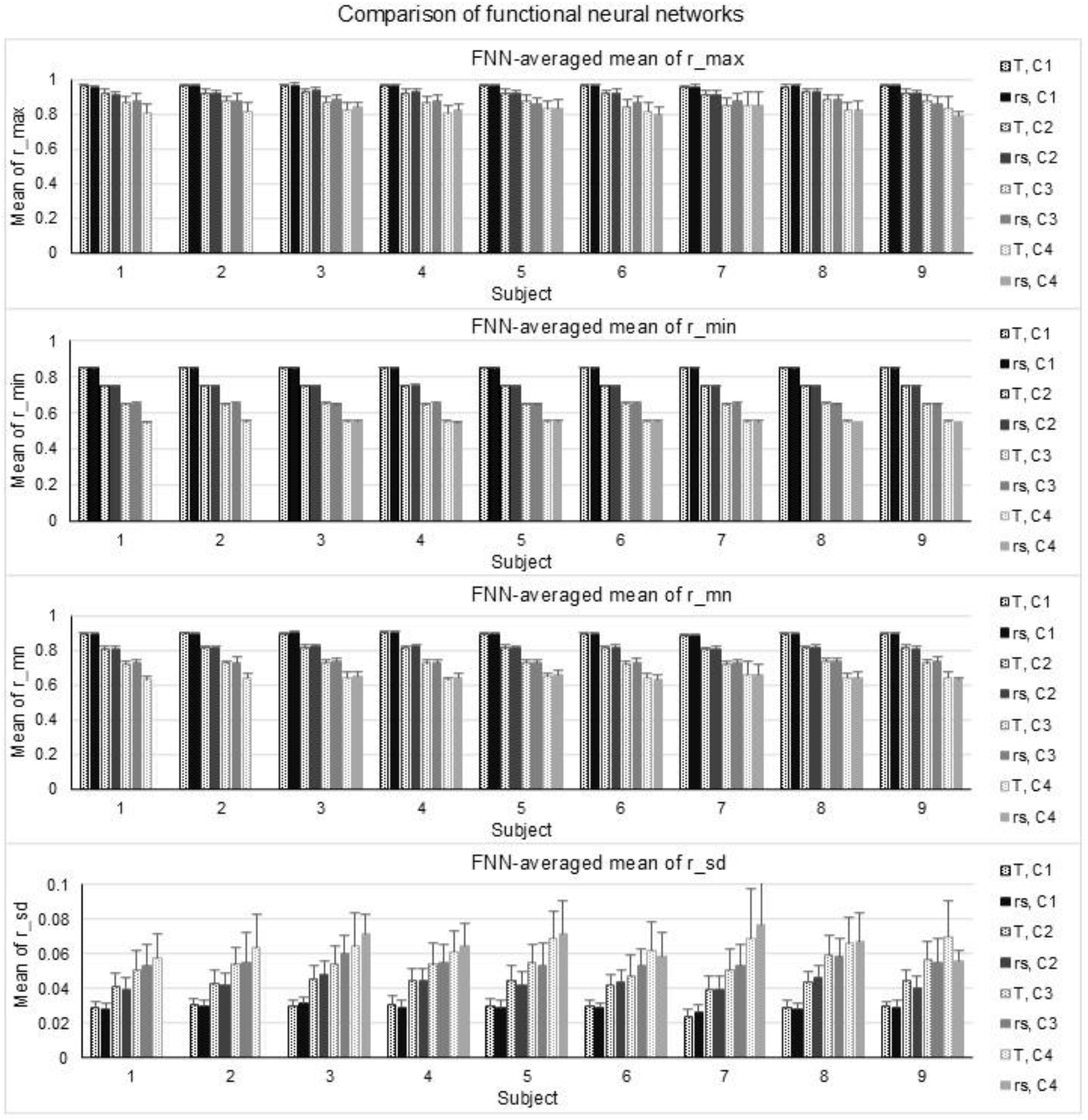
Characterization of the identified FNNs in each of the four categories for the two brain states and across the nine subjects. For each of the four parameters of a FNN (i.e., its maximum r, minimum r, mean r and sd), the mean value of that parameter was the one averaged over all FNNs in each brain state and each category for each subject. The error bar indicates its corresponding standard deviation (SD). T: task; and rs: resting-state.

### Quantification of the interaction between functional neural networks

The interaction between any two FNNs can be quantified by the correlation coefficient r of the neural activity between the two FNNs; the stronger the interaction, the larger the r value, and vice versa. For each FNN, the mean BOLD time signal averaged over all voxels reflects the dynamic neural activity across that FNN. For each paired FNNs, this r value reflects the functional relationship between the two FNNs. Functionally associated FNNs should yield large r values that show the statistically significant levels for their corresponding correlations. Fig. 3 illustrates the functional relationships between all FNNs in all four categories for each brain state for the representative subject, and these functional relationships for each of the other eight subjects are illustrated in Suppl. Fig. 2. These functional relationships between all FNNs showed a remarkable variation not only from state to state but also from subject to subject. Some FNNs were strongly associated with each other, some of them were negatively correlated with each other, and some of them were independent to each other as reflected in their corresponding small r values.

**Fig. 3.**
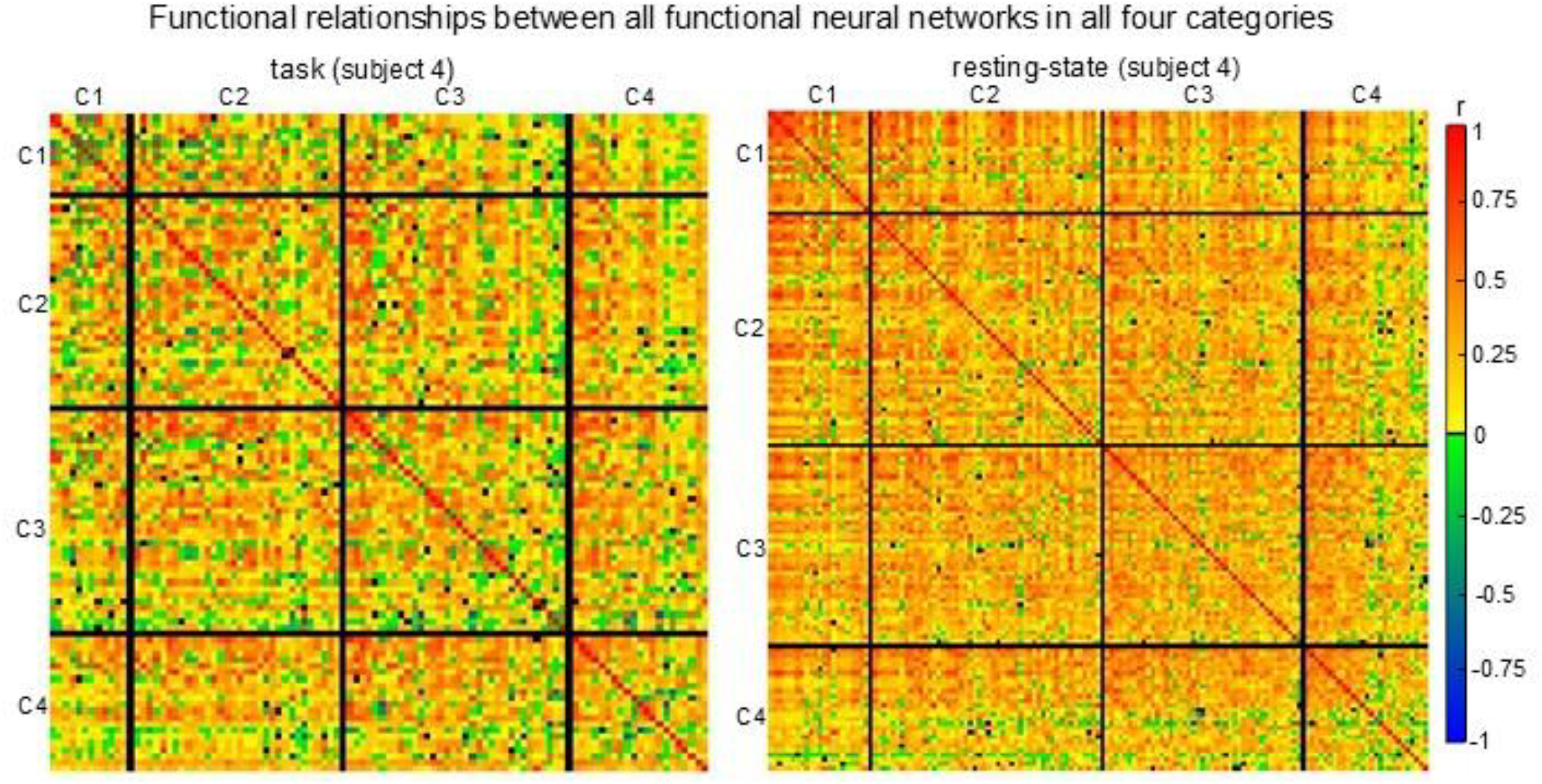
Illustration of the functional relationships between FNNs in all four categories for the two brain states for the representative subject. Each pixel denotes the correlation coefficient r of the neural activity between two FNNs and its color represents the value of r. Each black line separates the FNNs from one category to another.

### Symmetrically organized intrinsic neural activity

A FNN is unilateral if all voxels locate within one hemisphere of the brain; otherwise, it is bilateral. In each category, the total number of bilateral FNNs was substantially more than that of unilateral FNNs for every individual subject, and the group-mean number showed that the former is significantly larger than the latter (Fig. 4, left). A bilateral FNN is symmetric if it covers at least two anatomic areas that are positioned symmetrically across the left and right hemispheres of the brain; otherwise, it is non-symmetric. Again, in each category, the total number of symmetric FNNs was substantially more than that of non-symmetric FNNs for every individual subject, and the group-mean number also showed that the former is significantly larger than the latter (Fig. 4, right), revealing a symmetrically organized intrinsic neural activity.

**Fig. 4.**
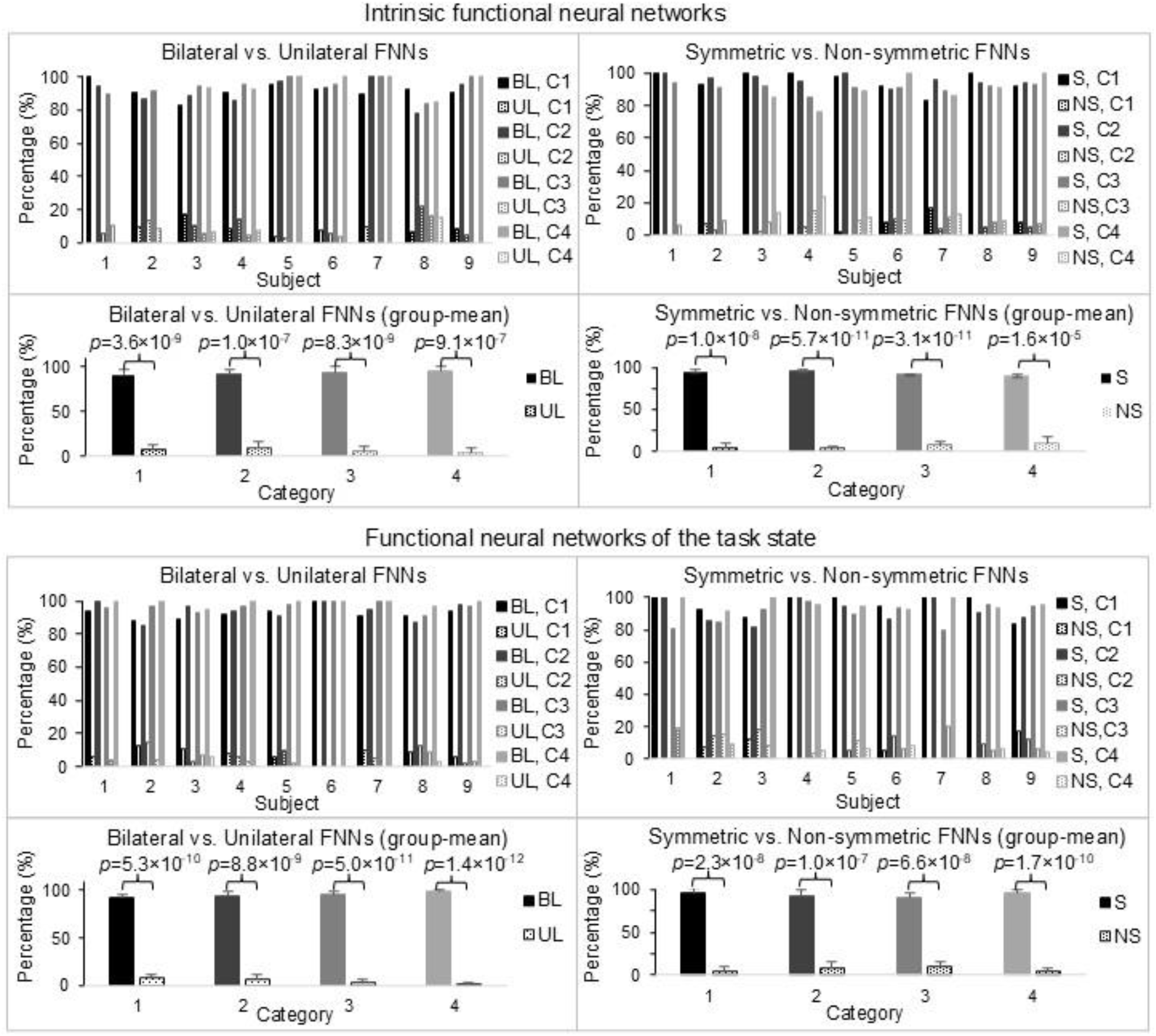
Comparisons of unilateral (UL) vs. bilateral (BL) and symmetric (S) vs. non-symmetric (NS) functional neural networks for both intrinsic neural activity and the neural activity when performing tasks. Each FNN was visually determined to be either UL or BL. For each category and each subject, the percentage of the total number of the UL or BL FNNs relative to the total number of the FNNs was computed, respectively. Similarly, each BL FNN was also visually determined to be either S or NS, and, for each category and each subject, the percentage of the total number of the S or NS FNNs relative to the total number of the BL FNNs was computed, respectively.

To illustrate this symmetrically organized intrinsic neural activity, we identified 18 symmetric FNNs for each individual subject (Fig. 5 and Suppl. Fig. 3, left three columns). For each symmetric FNN, the two anatomic areas covered by that FNN were symmetrically located across the left and right hemispheres of the brain for all subjects (Suppl. Figs. 4 and 5), and the high value of the correlation coefficient r (e.g., r>0.85) showed a synchronized neural activity between these two anatomic areas (Fig. 6). We computed the correlation coefficient r of the neural activity of each symmetric FNN with that of all other FNNs and then thresholded it with r>0.55 (*p*<1.0×10^−20^ for N=288) to determine its functionally connected FNNs for each subject (Fig. 5 and Suppl. Fig. 3, middle three columns). The right three columns in these two figures illustrate their corresponding group-mean functionally connected FNNs. For each symmetric FNN, its functionally connected FNNs and group-mean FNNs showed a symmetric neural activity across the left and right hemispheres of the brain, further demonstrating the symmetrically organized intrinsic neural activity at both individual and group levels.

**Fig. 5.**
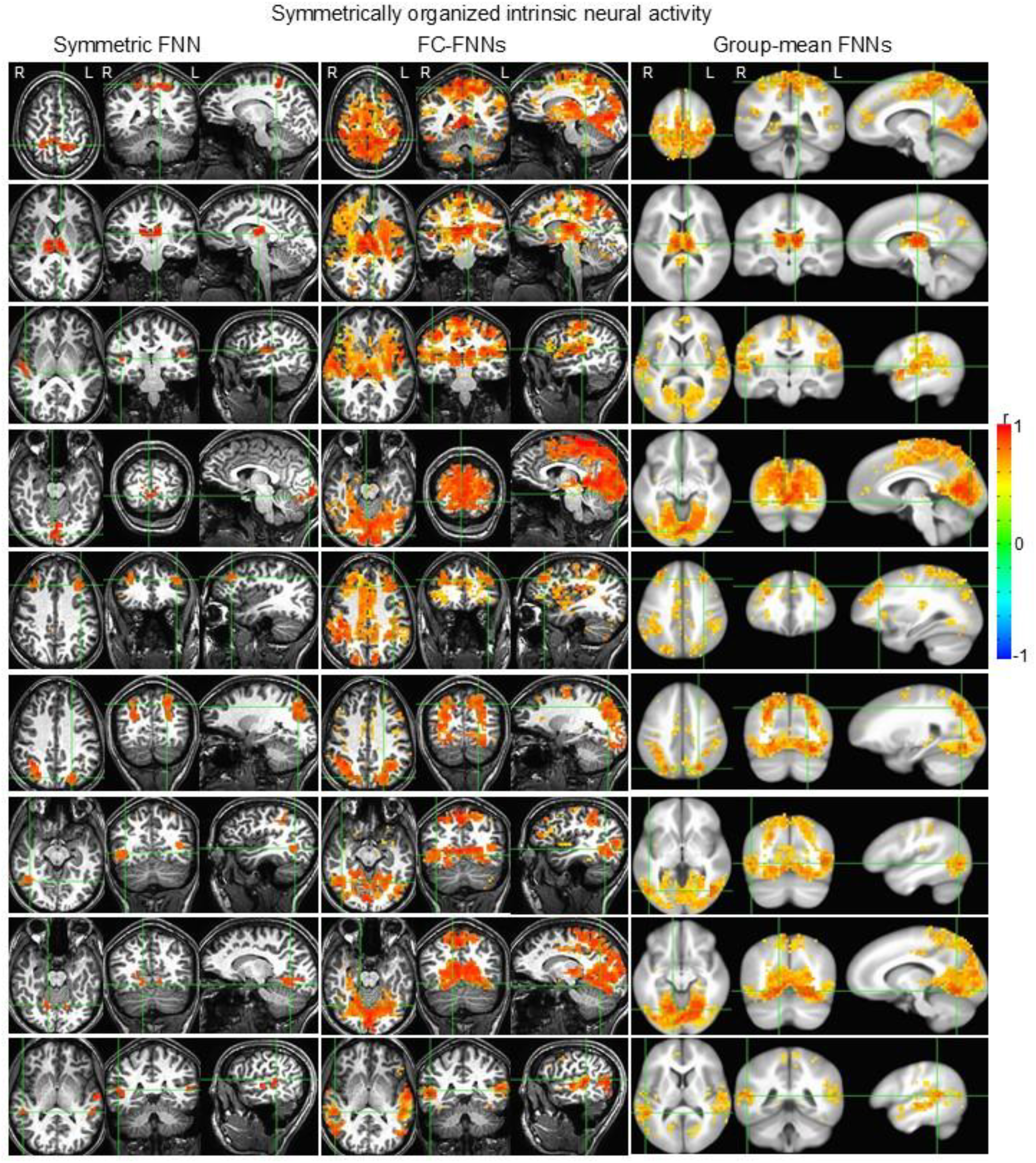
Illustration of nine identified symmetric FNNs (left three columns) with their corresponding functionally connected (FC) FNNs (middle three columns) for the representative subject and group-mean FC FNNs averaged over the nine subjects (right three columns). For each grouped axial, coronal and sagittal images in the left three columns, the two green lines in each image indicate the positions of the other two images, respectively. The crosspoint of these three lines indicates one of the two anatomic areas covered by that symmetric FNN.

**Fig. 6.**
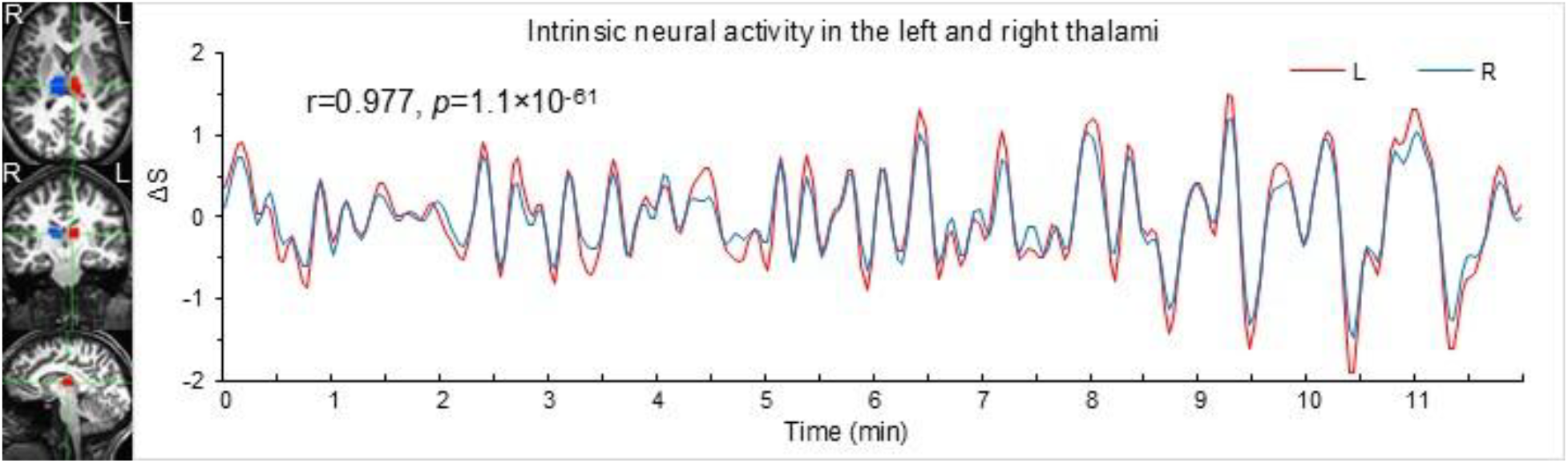
Illustration of the synchronized intrinsic neural activity between the left and right thalami for the representative subject (Fig. 5, the left three images in the 2^nd^ panel). The red region of interest (ROI) in the left three images indicates the left thalamus and the blue ROI indicates the right thalamus. For each thalamus, its neural activity is reflected in the mean BOLD time signal averaged over all voxels within that ROI. L: left; and R: right.

### Functional neural networks of the task state

Similarly to the intrinsic FNNs, the FNNs of the task state also showed the dominating bilateral and symmetric characteristics for each FNN category at both individual and group levels (Fig. 4, bottom panel). Using the identified 18 symmetric FNNs of the resting state as references, for each symmetric FNN we identified a similar symmetric FNN for the task state of each subject (Fig. 7 and Suppl. Fig. 6, left three columns). Similarly, we computed the correlation coefficient r of the neural activity of each symmetric FNN with that of all other FNNs and then thresholded it with r>0.55 (*p*<1.0×10^− 20^ for N=288) to determine its functionally connected FNNs for each subject (Fig. 7 and Suppl. Fig. 6, middle three columns). The right three columns in these two figures illustrate their corresponding group-mean functionally connected FNNs. For each symmetric FNN, similarly to the intrinsic neural activity, its functionally connected FNNs and group-mean FNNs also showed the characteristics of symmetric neural activity across the left and right hemispheres of the brain at both individual and group levels.

**Fig. 7.**
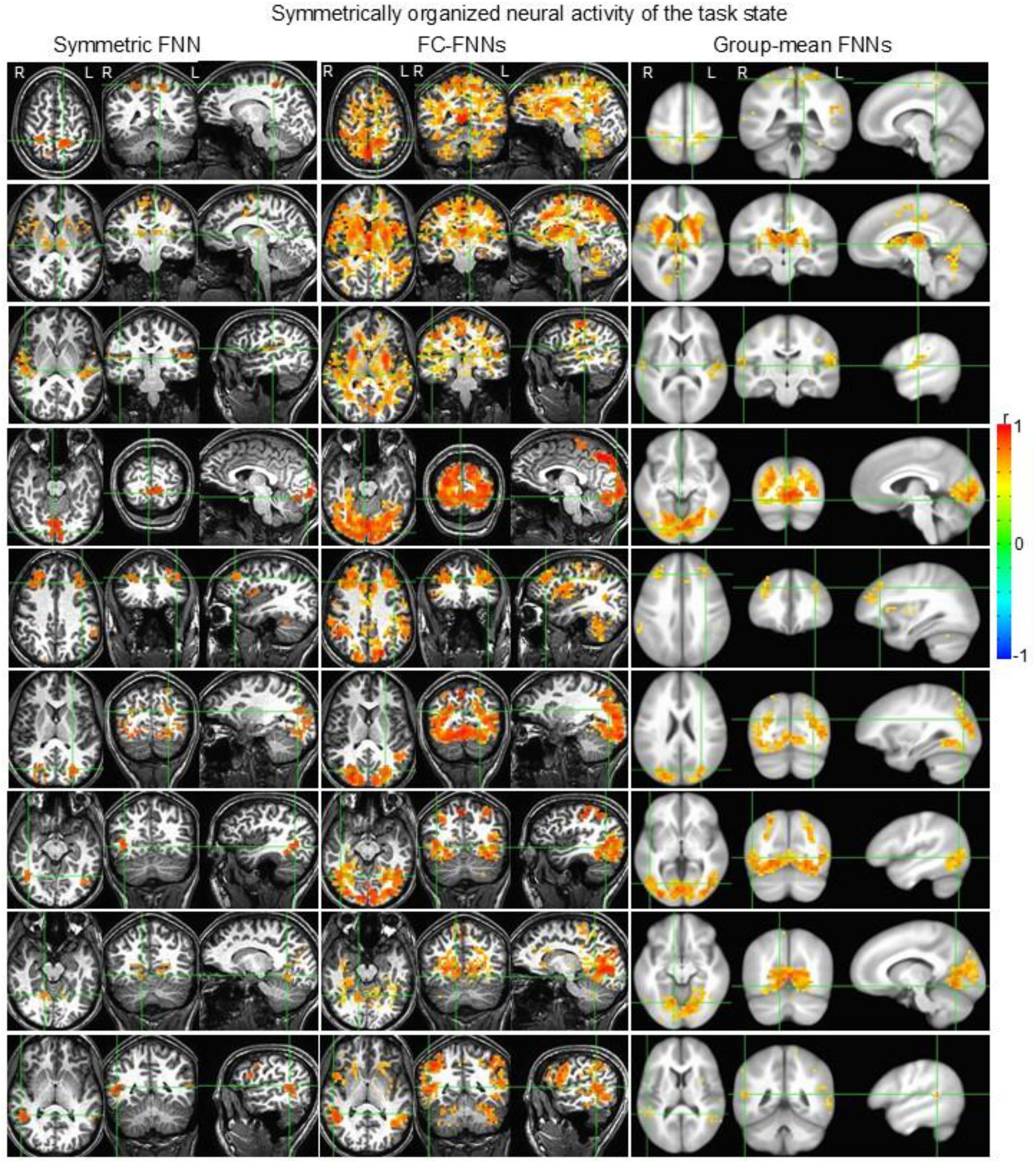
Illustration of nine identified symmetric FNNs (left three columns) with their corresponding functionally connected (FC) FNNs (middle three columns) for the representative subject and group-mean FC FNNs averaged over the nine subjects (right three columns). For each grouped axial, coronal and sagittal images in the left three columns, the two green lines in each image indicate the positions of the other two images, respectively. The crosspoint of these three lines indicates one of the two anatomic areas covered by that symmetric FNN.

The primary task of analyzing task-fMRI data is to identify task-evoked brain activated areas. The task paradigm consisted of three tasks of word-reading (WR), pattern-viewing (PV) and FT. Accordingly, there were seven task categories: (1) WR; (2) PV; (3) FT; (4) WR & PV; (5) WR & FT; (6) PV & FT; and (7) WR, PV & FT. For each task category, we computed the r of that category’s task-evoked ideal BOLD response with the time signal of each FNN, and then identified the task-associated FNNs with the threshold r value of 0.35 (*p*=2.9×10^−9^) for each subject (Suppl. Table 4). To validate this association, for each task category we computed the mean time signal averaged over all FNNs first for each subject and then averaged over all subjects. As expected, this mean time signal is significantly correlated with the task-evoked ideal BOLD response for each task category (maximum *p*=7.4×10^−33^), validating these task-associated FNNs for each task category (Suppl. Fig. 7). Substantial variations in the total number of the task-associated FNNs in each task category were observed from subject to subject (Supp. Table 4), indicating a varied task-evoked brain activation from subject to subject even when performing a simple task such as FT.

## Discussion and Conclusions

The identified FNNs and the quantification of the interaction between these FNNs characterize the whole-brain activity to a certain degree for each brain state and each individual subject. This identification and quantification are objective and automatic with no a priori knowledge, providing a means of objectively and holistically characterizing the whole brain activity of any brain functional state measured at large-scale systems level with either rs-or task-fMRI for each individual brain. Speaking generally, about 80% of the whole-brain volume are functionally connected as shown in these FNNs, regardless of whether the brain is at resting or performing tasks. These FNNs, however, vary substantially from brain state to state and from subject to subject, showing a remarkably variable whole-brain activity even when people perform the same task, which may characterize each individual brain activity.

The whole-brain activity and its FNNs are mainly symmetrically organized across the left and right hemispheres of the brain for both brain states (Fig. 4). This symmetrically organized activity is characterized by symmetric FNNs and their corresponding functionally connected FNNs for each symmetric FNN (Figs. 5 & 7 and Suppl. Figs. 3 & 6). For each symmetric FNN, the neural activity is synchronized between the two anatomic areas located symmetrically across the left and right hemispheres of the brain (Fig. 6). Note that a symmetric FNN does not imply that the size of the activated area is equal in these two anatomic areas or the size of the total activated areas is equal between the two hemispheres. A symmetric FNN may, however, covers multiple paired anatomic areas that are symmetrically located across the two hemispheres, and the neural activity is synchronized between these paired anatomic areas, showing an essential role in characterizing the whole-brain activity.

The whole-brain activity and its FNNs vary substantially not only from brain state to state but also from subject to subject (Fig. 1 and Suppl. Fig. 1). Each symmetric FNN and its functionally connected FNNs also vary substantially from subject to subject as illustrated in Suppl. Figs. 4 and 5. For each symmetric FNN, regardless of the brain state, its corresponding group-mean functionally connected FNNs reflects the commonality of that FNN across all subjects (Figs. 5 & 7 and Suppl. Figs. 3 & 6, right three columns). Its functionally connected FNNs, however, varies substantially from subject to subject as illustrated in Suppl. Figs. 4 and 5. For each subject, the difference of this functionally connected FNNs from its corresponding group-mean functionally connected FNNs reflects the individuality of that subject, providing a means of investigating the neural underpinnings of this individuality for each individual subject.

The brain’s operations are mainly intrinsic, involving the acquisition and maintenance of information for interpreting, responding to, and predicting environmental demands^6^. This spontaneous neural activity is mainly symmetrically organized across the left and right hemispheres of the brain for each individual brain. For the symmetric FNNs, the almost perfectly synchronized neural activity between each symmetrically located paired areas across the two hemispheres shows a constant communication between these paired areas (Fig. 6), indicating a functional coupling between these brain anatomic areas. The holistic characterization of the intrinsic neural activity across the entire brain with FNNs could be crucial for understanding the neural underpinnings of the brain’s operations at rest.

## Method and Materials

We extend our previous six studies^5,7-11^. This study analyzed the same fMRI data. It used the same subjects, same image acquisition, and same image preprocessing procedures. We briefly describe each paragraph. For more information, refer to our previous study^5^.

### Participants

Nine healthy subjects (4 female and 5 male, ages 21-55 years old) participated in the study. The Institutional Review Board at Michigan State University approved the study, and written informed consent was obtained from all subjects prior to the study. All methods were performed in accordance with the institution’s relevant guidelines and regulations.

### Image acquisition

Functional brain images were acquired on a GE 3.0 T clinical scanner with an 8-channel head coil using a gradient echo Echo-Planar-Imaging pulse sequence (TE/TR = 28/2500 ms, flip angle 80°, FOV 224 mm, matrix 64×64, slice thickness 3.5 mm, and spacing 0.0 mm). Thirty-eight axial slices to cover the entire brain were scanned, and the first three volume images were discarded. Each subject undertook a 12-min rs-fMRI scan and then a 12-min task-fMRI scan while performing three different tasks. Each scan yielded a total of 288 volume images (total time points N=288). For the rs-fMRI scan, the participants were instructed to close their eyes and try not to think of anything but remain awake during the scan. For the task-fMRI scan, each task was presented eight times, a total of 24 task trials, and the task presentation was interleaved. Each trial comprised a 6-s task period followed by a 24-s rest period, resulting in 12 volume images for each task trial. Task 1 was a word-reading paradigm: subjects silently read English words. Task 2 was a pattern-viewing paradigm: subjects viewed a black-and-white striped pattern. Task 3 was a visually cued FT paradigm: subjects tapped the five fingers of their right-hands as quick as possible in a random order. During the 24-s rest period, subjects were asked to focus their eyes on a small fixation mark at the screen center and try not to think of anything. After the task-fMRI scan, T1-weighted whole-brain MR images were also acquired using a 3D IR-SPGR pulse sequence.

### Image preprocessing

Image preprocessing of the functional images was performed using AFNI (analysis of functional neuro images) software^5,12^. It included removing spikes, slice-timing correction, motion correction, spatial filtering with a Gaussian kernel with a full-width-half-maximum of 4.0 mm, computing the mean volume image, bandpassing the signal intensity time courses to the range of 0.009–0.08 Hz, and computing the relative signal change (%) of the bandpassed signal intensity time courses. For each subject, based on the T1 and EPI images, we generated a mask to cover the entire brain for that subject. After these preprocessing steps, further image analysis was carried out using in-house developed Matlab-based software algorithms.

### Identification of FNN

This identification comprises 4 processes to identify FNNs in the four categories from C1 to C4 successively.

1. Identifying FNNs in C1

a. For a given voxel, finding a seed in which the correlation coefficient r of the BOLD time signal between two voxels is larger than 0.85 for pairwise combinations of all voxels within that seed and the total number of these voxels is equal to or larger than 10.
b. Testing all voxels to find all seeds.
c. Sorting the seeds according to the total number of voxels contained in each seed.
d. For the seed with the most voxels, computing the mean BOLD time signal averaged over these voxels. Computing the correlation coefficient r of this mean time signal with voxel’s time signal for all voxels across the entire brain and then identifying all voxels with r>0.85 to form a cluster. Computing the mean time signal of this cluster and using it to identify voxels with r>0.85 to update the cluster. Iterating this process until the cluster is stabilized, i.e., the voxels in the cluster remain unchanged.
e. If the total number of voxels in the cluster is equal to or larger than 50, these voxels constitute a FNN in C1. Then, excluding these voxels from further analysis.
f. Repeating procedures a) to e) to identify next FNN in C1.
g. Repeating procedures a) to f) to identify all FNNs in C1.

2. Identifying FNNs in C2, C3 and C4

Changing the r threshold value from 0.85 to 0.75 and repeating the same process to identify all FNNs in C2. Then, changing the threshold value to 0.65 and repeating the same process to identify all FNNs in C3. Finally, changing the value to 0.55 and repeating the same process to identify all FNNs in C4.

### Computation of the histogram of r distribution

For each brain state and each subject, we computed the correlation coefficient r of voxel’s time signal for pairwise combinations of all voxels that cover the entire brain. To compute the histogram of r distribution across the r range [-1, 1], we first divided the range to 200 equal intervals with interval size 0.01. Then, we counted the total number of paired voxels in each r interval and calculated its percentage (%) relative to the total number of all paired voxels within the entire brain, i.e., the sum of this histogram over the entire range [-1, 1] is equal to 100 (%).

### Task-induced ideal BOLD response and image group analysis

For each task category, the task-induced ideal BOLD response was generated by convolving the temporal paradigm of that task with a hemodynamic response function, using the 3dDeconvolve program in AFNI with the convolution kernel SPMG3. For image group analysis, each subject’s brain images were converted to a standard template space (icbm452, an averaged volume of 452 normal brains) using AFNI.

## Acknowledgements

This work was supported by the Michigan State University Radiology Pilot Scan Program.

## Author Contributions

The sole author is fully responsible for the paper.

## Declaration of generative AI in scientific writing

No generative AI and AI-assisted technologies were used in writing this manuscript.

## Informed consent statement

The Institutional Review Board at Michigan State University approved the study, and all methods were performed in accordance with the institution’s relevant guidelines and regulations. Written informed consent was obtained from all subjects prior to the study.

## Data availability

Both the original and processed fMRI images plus final research data related to this publication will be available to share upon request from the corresponding author with a legitimate reason such as to validate the reported findings or to conduct a new analysis.

