## Supplemental Figures for "Identification of functional neural networks of human brains with fMRI"

Jie Huang

Department of Radiology

Michigan State University

East Lansing, MI, USA

Suppl. Table 1. Functional neural networks in category 2 determined for both brain states for the representative subject. #: number; r_max: maximum r; r_min: minimum r; mn: mean r value averaged over all voxels within each FNN; and sd: the corresponding standard deviation of these r values.

| Task | | | | | | resting-state | | | | | |
|---|---|---|---|---|---|---|---|---|---|---|---|
| | | Pearson Correlation Coefficient r | | | | | | Pearson Correlation Coefficient r | | | |
| FNN | # of voxels | r_max | r_min | r_mn | r_sd | FNN | # of voxels | r_max | r_min | r_mn | r_sd |
| 1 | 449 | 0.917 | 0.751 | 0.801 | 0.036 | 1 | 407 | 0.908 | 0.750 | 0.799 | 0.036 |
| 2 | 441 | 0.885 | 0.750 | 0.799 | 0.033 | 2 | 404 | 0.904 | 0.751 | 0.799 | 0.033 |
| 3 | 378 | 0.939 | 0.751 | 0.802 | 0.040 | 3 | 315 | 0.942 | 0.750 | 0.824 | 0.046 |
| 4 | 273 | 0.929 | 0.750 | 0.804 | 0.037 | 4 | 246 | 0.897 | 0.751 | 0.803 | 0.036 |
| 5 | 200 | 0.935 | 0.752 | 0.806 | 0.039 | 5 | 185 | 0.892 | 0.750 | 0.801 | 0.034 |
| 6 | 191 | 0.905 | 0.751 | 0.800 | 0.036 | 6 | 171 | 0.917 | 0.751 | 0.809 | 0.041 |
| 7 | 186 | 0.929 | 0.754 | 0.804 | 0.044 | 7 | 169 | 0.916 | 0.751 | 0.812 | 0.042 |
| 8 | 175 | 0.885 | 0.753 | 0.804 | 0.032 | 8 | 152 | 0.935 | 0.752 | 0.818 | 0.047 |
| 9 | 167 | 0.929 | 0.751 | 0.801 | 0.039 | 9 | 129 | 0.942 | 0.756 | 0.831 | 0.043 |
| 10 | 159 | 0.947 | 0.752 | 0.818 | 0.051 | 10 | 127 | 0.945 | 0.750 | 0.827 | 0.054 |
| 11 | 143 | 0.920 | 0.750 | 0.822 | 0.048 | 11 | 120 | 0.896 | 0.753 | 0.802 | 0.037 |
| 12 | 143 | 0.920 | 0.751 | 0.815 | 0.044 | 12 | 119 | 0.914 | 0.750 | 0.815 | 0.042 |
| 13 | 143 | 0.923 | 0.752 | 0.810 | 0.045 | 13 | 119 | 0.915 | 0.754 | 0.805 | 0.038 |
| 14 | 136 | 0.950 | 0.751 | 0.820 | 0.051 | 14 | 116 | 0.931 | 0.759 | 0.819 | 0.040 |
| 15 | 120 | 0.912 | 0.753 | 0.809 | 0.039 | 15 | 111 | 0.920 | 0.753 | 0.813 | 0.042 |
| 16 | 110 | 0.923 | 0.752 | 0.813 | 0.040 | 16 | 109 | 0.933 | 0.751 | 0.823 | 0.045 |
| 17 | 99 | 0.933 | 0.756 | 0.821 | 0.048 | 17 | 108 | 0.931 | 0.751 | 0.801 | 0.040 |
| 18 | 97 | 0.927 | 0.755 | 0.820 | 0.048 | 18 | 102 | 0.919 | 0.757 | 0.823 | 0.049 |
| 19 | 92 | 0.931 | 0.753 | 0.815 | 0.044 | 19 | 101 | 0.919 | 0.752 | 0.809 | 0.041 |
| 20 | 89 | 0.915 | 0.752 | 0.826 | 0.048 | 20 | 96 | 0.912 | 0.755 | 0.819 | 0.044 |
| 21 | 89 | 0.937 | 0.753 | 0.811 | 0.047 | 21 | 96 | 0.905 | 0.751 | 0.811 | 0.042 |

| | | | | | | | | | | | |
|---|---|---|---|---|---|---|---|---|---|---|---|
| 22 | 88 | 0.908 | 0.755 | 0.813 | 0.038 | 22 | 94 | 0.921 | 0.751 | 0.822 | 0.045 |
| 23 | 86 | 0.950 | 0.755 | 0.814 | 0.048 | 23 | 90 | 0.921 | 0.752 | 0.823 | 0.043 |
| 24 | 79 | 0.906 | 0.752 | 0.817 | 0.043 | 24 | 87 | 0.923 | 0.751 | 0.808 | 0.039 |
| 25 | 78 | 0.954 | 0.750 | 0.835 | 0.054 | 25 | 86 | 0.938 | 0.757 | 0.829 | 0.041 |
| 26 | 77 | 0.924 | 0.751 | 0.816 | 0.048 | 26 | 85 | 0.955 | 0.760 | 0.837 | 0.053 |
| 27 | 72 | 0.925 | 0.751 | 0.822 | 0.049 | 27 | 81 | 0.904 | 0.753 | 0.819 | 0.040 |
| 28 | 72 | 0.893 | 0.751 | 0.812 | 0.040 | 28 | 78 | 0.914 | 0.755 | 0.815 | 0.043 |
| 29 | 71 | 0.934 | 0.761 | 0.839 | 0.050 | 29 | 78 | 0.900 | 0.752 | 0.811 | 0.039 |
| 30 | 68 | 0.938 | 0.755 | 0.821 | 0.048 | 30 | 73 | 0.957 | 0.753 | 0.830 | 0.054 |
| 31 | 58 | 0.954 | 0.750 | 0.837 | 0.063 | 31 | 72 | 0.910 | 0.760 | 0.823 | 0.041 |
| 32 | 52 | 0.951 | 0.754 | 0.836 | 0.055 | 32 | 69 | 0.901 | 0.751 | 0.810 | 0.039 |
| | | | | | | 33 | 68 | 0.970 | 0.758 | 0.857 | 0.060 |
| | | | | | | 34 | 68 | 0.933 | 0.755 | 0.814 | 0.043 |
| | | | | | | 35 | 67 | 0.920 | 0.751 | 0.811 | 0.044 |
| | | | | | | 36 | 67 | 0.932 | 0.755 | 0.843 | 0.052 |
| | | | | | | 37 | 67 | 0.935 | 0.751 | 0.813 | 0.052 |
| | | | | | | 38 | 65 | 0.918 | 0.756 | 0.813 | 0.041 |
| | | | | | | 39 | 64 | 0.956 | 0.752 | 0.829 | 0.060 |
| | | | | | | 40 | 62 | 0.937 | 0.757 | 0.829 | 0.052 |
| | | | | | | 41 | 61 | 0.918 | 0.757 | 0.827 | 0.047 |
| | | | | | | 42 | 60 | 0.938 | 0.754 | 0.828 | 0.051 |
| | | | | | | 43 | 59 | 0.955 | 0.759 | 0.841 | 0.056 |
| | | | | | | 44 | 58 | 0.908 | 0.751 | 0.808 | 0.041 |
| | | | | | | 45 | 57 | 0.913 | 0.753 | 0.815 | 0.039 |
| | | | | | | 46 | 56 | 0.952 | 0.760 | 0.835 | 0.047 |
| | | | | | | 47 | 55 | 0.949 | 0.753 | 0.840 | 0.048 |
| | | | | | | 48 | 54 | 0.952 | 0.752 | 0.839 | 0.049 |
| | | | | | | 49 | 52 | 0.931 | 0.760 | 0.844 | 0.054 |
| | | | | | | 50 | 50 | 0.960 | 0.755 | 0.842 | 0.054 |
| mean | 146 | 0.926 | 0.752 | 0.815 | 0.044 | mean | 110 | 0.926 | 0.754 | 0.820 | 0.045 |
| SD | 104 | 0.018 | 0.002 | 0.011 | 0.007 | SD | 79 | 0.019 | 0.003 | 0.014 | 0.007 |

Suppl. Table 2. Functional neural networks in category 3 determined for both brain states for the representative subject. #: number; r_max: maximum r; r_min: minimum r; mn: mean r value averaged over all voxels within each FNN; and sd: the corresponding standard deviation of these r values.

| Task | | | | | | resting-state | | |
|---|---|---|---|---|---|---|---|---|
| | | Pearson Correlation Coefficient r | | | | | | Pearson Correlation Coefficient r |

| FNN | # of voxels | r_max | r_min | r_mn | r_sd | FNN | # of voxels | r_max | r_min | r_mn | r_sd |
|---|---|---|---|---|---|---|---|---|---|---|---|
| 1 | 882 | 0.825 | 0.650 | 0.698 | 0.036 | 1 | 363 | 0.839 | 0.651 | 0.714 | 0.043 |
| 2 | 649 | 0.856 | 0.650 | 0.702 | 0.038 | 2 | 356 | 0.842 | 0.650 | 0.700 | 0.040 |
| 3 | 466 | 0.799 | 0.651 | 0.696 | 0.032 | 3 | 340 | 0.819 | 0.651 | 0.704 | 0.037 |
| 4 | 317 | 0.891 | 0.651 | 0.720 | 0.050 | 4 | 290 | 0.883 | 0.651 | 0.724 | 0.055 |
| 5 | 301 | 0.844 | 0.651 | 0.717 | 0.047 | 5 | 241 | 0.865 | 0.650 | 0.723 | 0.050 |
| 6 | 272 | 0.891 | 0.651 | 0.721 | 0.053 | 6 | 240 | 0.906 | 0.651 | 0.714 | 0.054 |
| 7 | 243 | 0.847 | 0.650 | 0.716 | 0.045 | 7 | 234 | 0.886 | 0.653 | 0.720 | 0.054 |
| 8 | 227 | 0.903 | 0.651 | 0.719 | 0.052 | 8 | 217 | 0.846 | 0.650 | 0.716 | 0.044 |
| 9 | 221 | 0.874 | 0.652 | 0.710 | 0.046 | 9 | 181 | 0.908 | 0.651 | 0.726 | 0.058 |
| 10 | 211 | 0.875 | 0.651 | 0.713 | 0.050 | 10 | 167 | 0.886 | 0.650 | 0.717 | 0.053 |
| 11 | 155 | 0.852 | 0.651 | 0.722 | 0.048 | 11 | 167 | 0.839 | 0.650 | 0.710 | 0.042 |
| 12 | 145 | 0.913 | 0.651 | 0.723 | 0.058 | 12 | 163 | 0.919 | 0.652 | 0.726 | 0.054 |
| 13 | 142 | 0.819 | 0.650 | 0.716 | 0.039 | 13 | 156 | 0.882 | 0.650 | 0.729 | 0.061 |
| 14 | 138 | 0.894 | 0.652 | 0.726 | 0.059 | 14 | 146 | 0.805 | 0.652 | 0.705 | 0.036 |
| 15 | 137 | 0.871 | 0.655 | 0.726 | 0.055 | 15 | 140 | 0.831 | 0.656 | 0.726 | 0.047 |
| 16 | 131 | 0.839 | 0.652 | 0.711 | 0.045 | 16 | 138 | 0.840 | 0.650 | 0.723 | 0.047 |
| 17 | 127 | 0.883 | 0.655 | 0.734 | 0.057 | 17 | 137 | 0.877 | 0.654 | 0.755 | 0.057 |
| 18 | 127 | 0.843 | 0.654 | 0.731 | 0.053 | 18 | 136 | 0.919 | 0.653 | 0.740 | 0.062 |
| 19 | 125 | 0.888 | 0.650 | 0.727 | 0.060 | 19 | 134 | 0.843 | 0.654 | 0.712 | 0.043 |
| 20 | 122 | 0.853 | 0.651 | 0.718 | 0.048 | 20 | 121 | 0.826 | 0.653 | 0.713 | 0.043 |
| 21 | 113 | 0.872 | 0.651 | 0.723 | 0.057 | 21 | 119 | 0.909 | 0.651 | 0.730 | 0.059 |
| 22 | 113 | 0.839 | 0.652 | 0.715 | 0.046 | 22 | 114 | 0.892 | 0.651 | 0.725 | 0.056 |
| 23 | 101 | 0.849 | 0.658 | 0.722 | 0.048 | 23 | 113 | 0.873 | 0.652 | 0.731 | 0.052 |
| 24 | 101 | 0.842 | 0.654 | 0.717 | 0.047 | 24 | 108 | 0.894 | 0.652 | 0.738 | 0.059 |
| 25 | 84 | 0.904 | 0.651 | 0.750 | 0.064 | 25 | 107 | 0.829 | 0.652 | 0.724 | 0.045 |
| 26 | 81 | 0.864 | 0.651 | 0.728 | 0.052 | 26 | 102 | 0.878 | 0.652 | 0.733 | 0.060 |
| 27 | 81 | 0.902 | 0.654 | 0.739 | 0.072 | 27 | 101 | 0.854 | 0.654 | 0.711 | 0.046 |
| 28 | 80 | 0.913 | 0.656 | 0.751 | 0.067 | 28 | 95 | 0.924 | 0.661 | 0.752 | 0.063 |
| 29 | 67 | 0.915 | 0.654 | 0.752 | 0.066 | 29 | 93 | 0.876 | 0.662 | 0.741 | 0.055 |
| 30 | 65 | 0.917 | 0.651 | 0.771 | 0.066 | 30 | 93 | 0.879 | 0.653 | 0.736 | 0.054 |
| 31 | 61 | 0.863 | 0.659 | 0.730 | 0.055 | 31 | 91 | 0.883 | 0.652 | 0.733 | 0.061 |
| 32 | 60 | 0.909 | 0.650 | 0.733 | 0.058 | 32 | 87 | 0.845 | 0.650 | 0.723 | 0.043 |
| 33 | 55 | 0.934 | 0.652 | 0.770 | 0.086 | 33 | 77 | 0.904 | 0.652 | 0.739 | 0.065 |
| 34 | 50 | 0.931 | 0.660 | 0.784 | 0.086 | 34 | 77 | 0.899 | 0.659 | 0.745 | 0.058 |
| | | | | | | 35 | 76 | 0.902 | 0.656 | 0.736 | 0.066 |
| | | | | | | 36 | 73 | 0.892 | 0.655 | 0.754 | 0.065 |
| | | | | | | 37 | 73 | 0.888 | 0.656 | 0.757 | 0.054 |
| | | | | | | 38 | 66 | 0.884 | 0.656 | 0.750 | 0.063 |
| | | | | | | 39 | 61 | 0.901 | 0.651 | 0.747 | 0.067 |
| | | | | | | 40 | 56 | 0.901 | 0.655 | 0.740 | 0.063 |
| | | | | | | 41 | 54 | 0.937 | 0.651 | 0.772 | 0.078 |
| | | | | | | 42 | 53 | 0.917 | 0.656 | 0.740 | 0.069 |
| | | | | | | 43 | 51 | 0.916 | 0.656 | 0.741 | 0.081 |

| | | | | | | | | | | | |
|---|---|---|---|---|---|---|---|---|---|---|---|
| mean | 184 | 0.834 | 0.652 | 0.727 | 0.054 | mean | 140 | 0.878 | 0.653 | 0.730 | 0.055 |
| SD | 175 | 0.034 | 0.003 | 0.020 | 0.012 | SD | 82 | 0.032 | 0.003 | 0.016 | 0.010 |

Suppl. Table 3. Functional neural networks in category 4 determined for both brain states for the representative subject. #: number; r_max: maximum r; r_min: minimum r; mn: mean r value averaged over all voxels within each FNN; and sd: the corresponding standard deviation of these r values.

| Task | | | | | | resting-state | | | | | |
|---|---|---|---|---|---|---|---|---|---|---|---|
| | | Pearson Correlation Coefficient r | | | | | | Pearson Correlation Coefficient r | | | |
| FNN | # of voxels | r_max | r_min | r_mn | r_sd | FNN | # of voxels | r_max | r_min | r_mn | r_sd |
| 1 | 1143 | 0.712 | 0.550 | 0.602 | 0.036 | 1 | 479 | 0.769 | 0.551 | 0.607 | 0.044 |
| 2 | 555 | 0.775 | 0.552 | 0.617 | 0.051 | 2 | 403 | 0.836 | 0.550 | 0.633 | 0.064 |
| 3 | 500 | 0.829 | 0.550 | 0.619 | 0.056 | 3 | 378 | 0.795 | 0.550 | 0.613 | 0.050 |
| 4 | 406 | 0.773 | 0.550 | 0.615 | 0.049 | 4 | 276 | 0.836 | 0.551 | 0.642 | 0.064 |
| 5 | 291 | 0.785 | 0.551 | 0.616 | 0.051 | 5 | 228 | 0.739 | 0.552 | 0.614 | 0.043 |
| 6 | 243 | 0.744 | 0.550 | 0.615 | 0.045 | 6 | 216 | 0.783 | 0.552 | 0.619 | 0.052 |
| 7 | 221 | 0.787 | 0.551 | 0.633 | 0.055 | 7 | 189 | 0.888 | 0.553 | 0.631 | 0.066 |
| 8 | 216 | 0.825 | 0.551 | 0.621 | 0.057 | 8 | 184 | 0.806 | 0.551 | 0.636 | 0.067 |
| 9 | 164 | 0.838 | 0.551 | 0.636 | 0.059 | 9 | 175 | 0.800 | 0.551 | 0.630 | 0.060 |
| 10 | 150 | 0.846 | 0.554 | 0.639 | 0.069 | 10 | 167 | 0.785 | 0.551 | 0.629 | 0.053 |
| 11 | 148 | 0.838 | 0.551 | 0.630 | 0.064 | 11 | 159 | 0.839 | 0.550 | 0.651 | 0.064 |
| 12 | 127 | 0.854 | 0.552 | 0.630 | 0.071 | 12 | 158 | 0.832 | 0.551 | 0.639 | 0.068 |
| 13 | 126 | 0.737 | 0.551 | 0.618 | 0.048 | 13 | 151 | 0.806 | 0.552 | 0.650 | 0.057 |
| 14 | 119 | 0.801 | 0.550 | 0.621 | 0.052 | 14 | 134 | 0.846 | 0.560 | 0.642 | 0.067 |
| 15 | 117 | 0.879 | 0.550 | 0.648 | 0.079 | 15 | 129 | 0.834 | 0.553 | 0.634 | 0.062 |
| 16 | 116 | 0.885 | 0.554 | 0.648 | 0.085 | 16 | 115 | 0.869 | 0.550 | 0.658 | 0.082 |
| 17 | 109 | 0.816 | 0.553 | 0.643 | 0.062 | 17 | 109 | 0.814 | 0.554 | 0.633 | 0.061 |
| 18 | 105 | 0.810 | 0.550 | 0.641 | 0.066 | 18 | 107 | 0.793 | 0.553 | 0.634 | 0.054 |
| 19 | 88 | 0.843 | 0.552 | 0.643 | 0.066 | 19 | 99 | 0.863 | 0.555 | 0.670 | 0.073 |
| 20 | 72 | 0.824 | 0.564 | 0.649 | 0.074 | 20 | 99 | 0.814 | 0.552 | 0.659 | 0.071 |
| 21 | 69 | 0.818 | 0.561 | 0.661 | 0.078 | 21 | 78 | 0.825 | 0.551 | 0.657 | 0.074 |
| | | | | | | 22 | 77 | 0.798 | 0.564 | 0.651 | 0.060 |
| | | | | | | 23 | 76 | 0.814 | 0.558 | 0.649 | 0.061 |
| | | | | | | 24 | 74 | 0.874 | 0.557 | 0.689 | 0.099 |
| | | | | | | 25 | 60 | 0.868 | 0.550 | 0.658 | 0.071 |
| | | | | | | 26 | 59 | 0.816 | 0.556 | 0.689 | 0.082 |
| | | | | | | 27 | 56 | 0.866 | 0.556 | 0.672 | 0.081 |
| mean | 242 | 0.810 | 0.552 | 0.631 | 0.061 | mean | 164 | 0.822 | 0.553 | 0.644 | 0.065 |
| SD | 247 | 0.045 | 0.004 | 0.015 | 0.012 | SD | 109 | 0.035 | 0.003 | 0.021 | 0.012 |

Suppl. Table 4. Number of task-associated FNNs in each task category.

| | Number of FNNs | | | | | | | |
|---|---|---|---|---|---|---|---|---|
| Subject | WR | PV | FT | WR_PV | WR_FT | PV_FT | WR_PV_FT | Total |
| 1 | 8 | 11 | 12 | 14 | 15 | 3 | 28 | 91 |
| 2 | 12 | 8 | 29 | 19 | 33 | 22 | 29 | 152 |
| 3 | 7 | 8 | 11 | 11 | 16 | 15 | 18 | 86 |
| 4 | 4 | 10 | 6 | 13 | 1 | 8 | 6 | 48 |
| 5 | 2 | 13 | 34 | 9 | 15 | 34 | 20 | 127 |
| 6 | 8 | 7 | 22 | 13 | 30 | 28 | 34 | 142 |
| 7 | 2 | 2 | 4 | 4 | 4 | 2 | 1 | 19 |
| 8 | 5 | 11 | 18 | 9 | 18 | 17 | 12 | 90 |
| 9 | 33 | 15 | 14 | 30 | 30 | 14 | 30 | 166 |

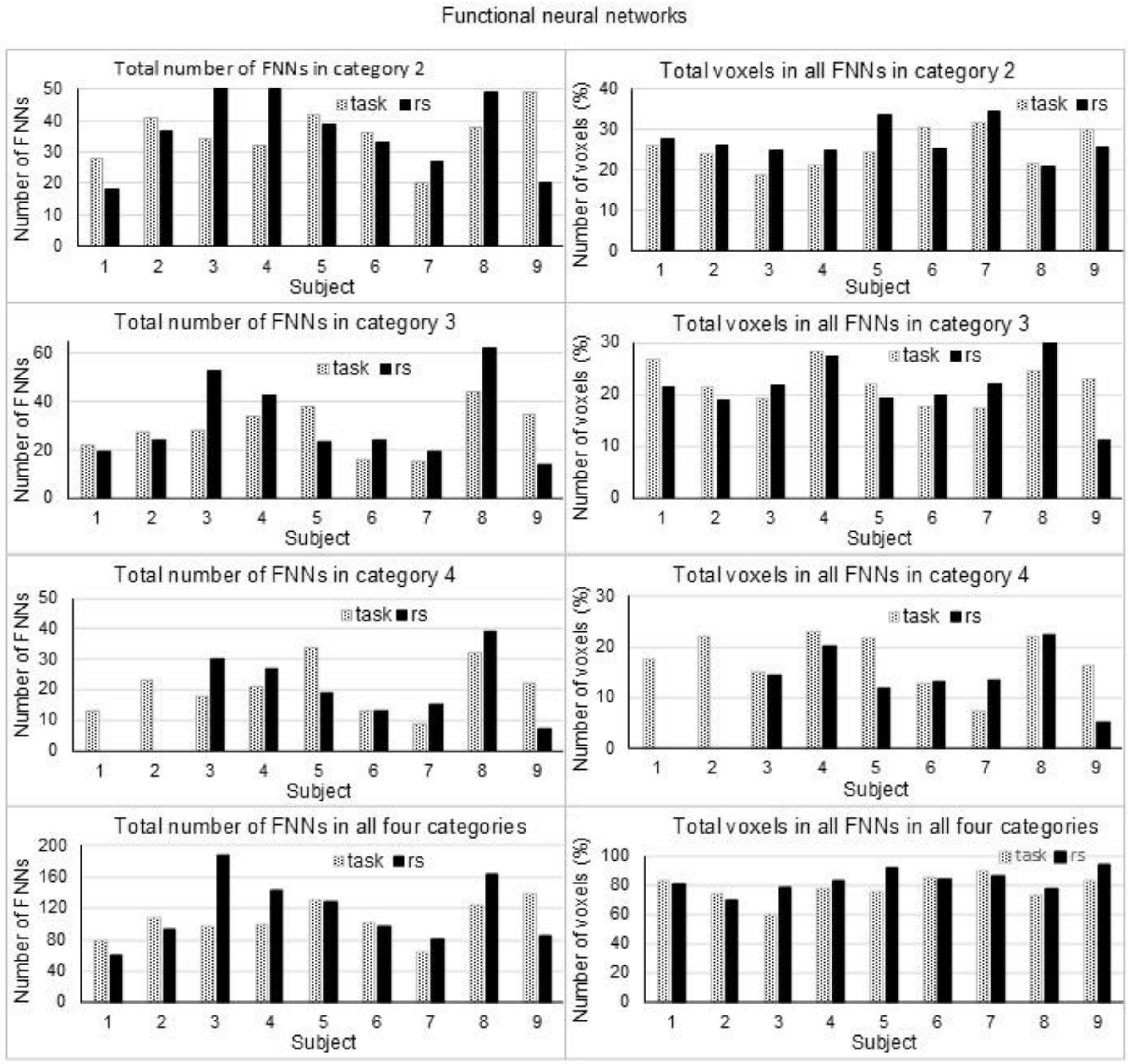

Suppl. Fig. 1 Comparisons of the identified FNNs in each of the three categories from 2 to 4 and in all four categories between the two brain states and across the nine subjects. The left plots illustrate the substantially varied total number of FNNs between the two brain states and across the nine subjects for all categories. The right plots illustrate their corresponding percentage of total number of voxels in these FNNs relative to the total number of voxels of the entire brain.

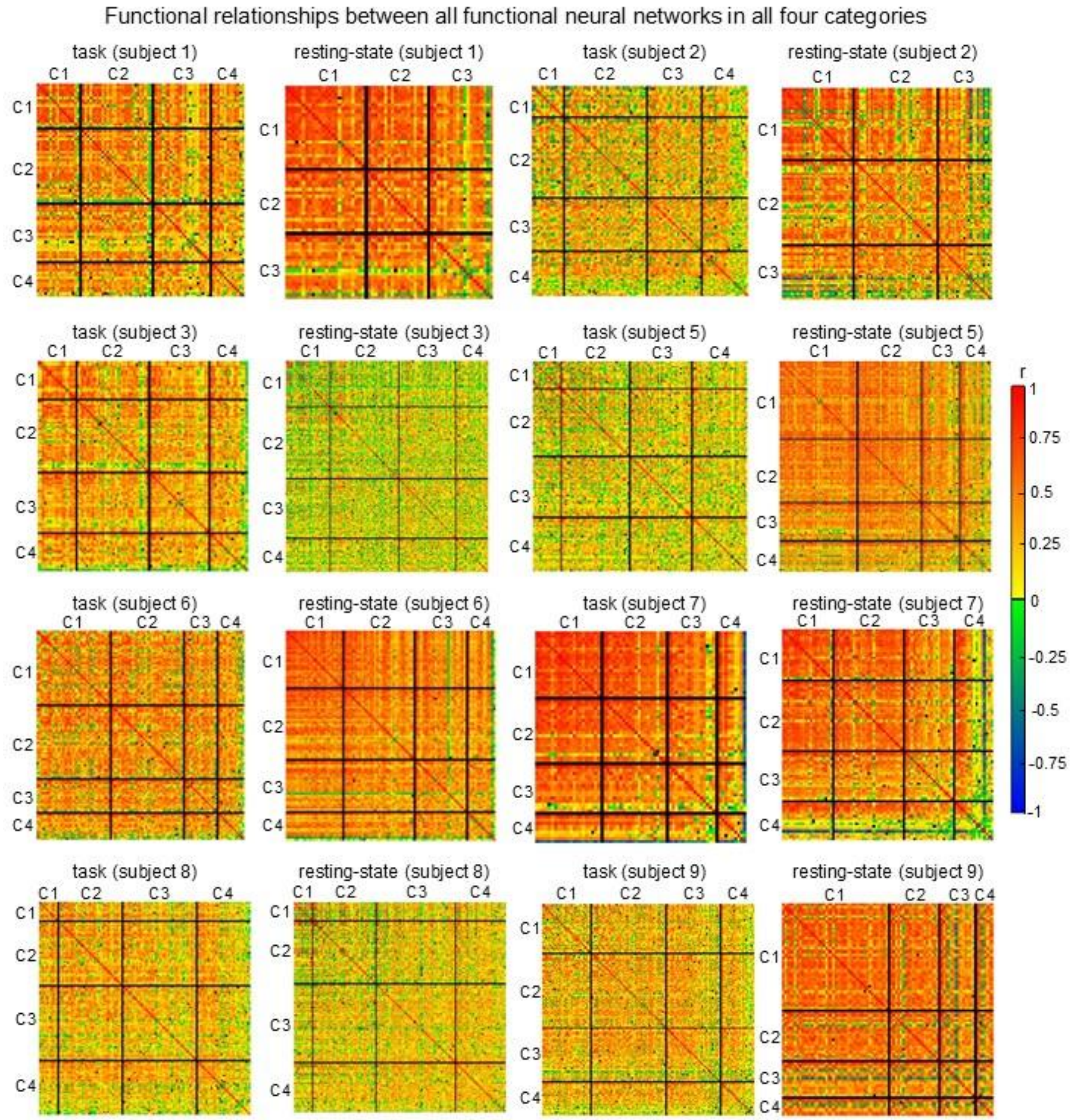

Suppl. Fig. 2 Illustration of the functional relationships between FNNs in all four categories for the two brain states for each of the rest eight subjects. Each pixel denotes the correlation coefficient r of the neural activity between two FNNs and its color represents the value of r. Each black line separates the FNNs from one category to another.

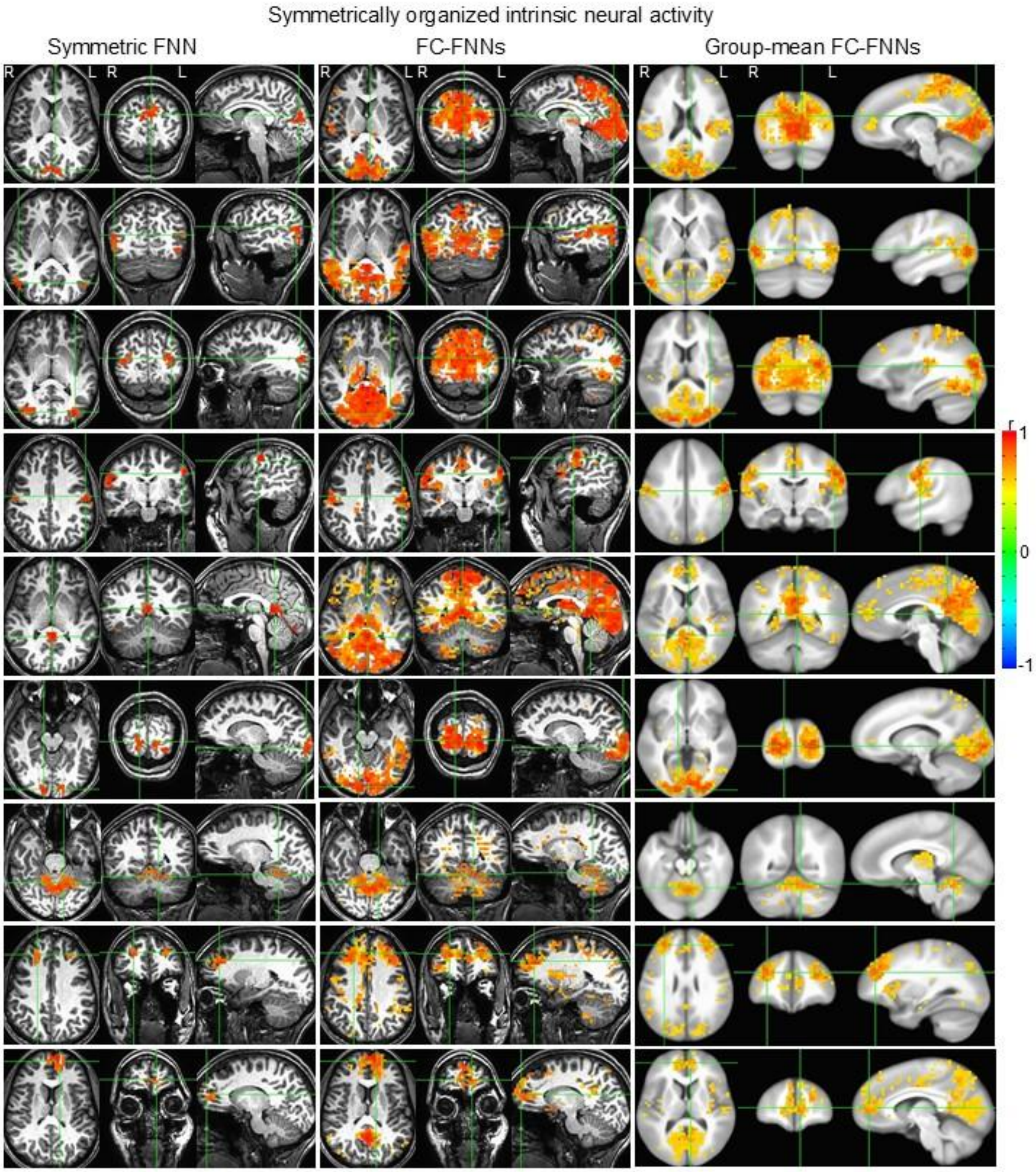

Suppl. Fig. 3 Illustration of the rest nine identified symmetric FNNs (left three columns) with their corresponding functionally connected (FC) FNNs (middle three columns) for the representative subject and group-mean FC FNNs averaged over the nine subjects (right three columns). For each grouped axial, coronal and sagittal images in the left three columns, the two green lines in each image indicate the positions of the other two images, respectively. The crosspoint of these three lines indicates one of the two anatomic areas covered by that symmetric FNN.

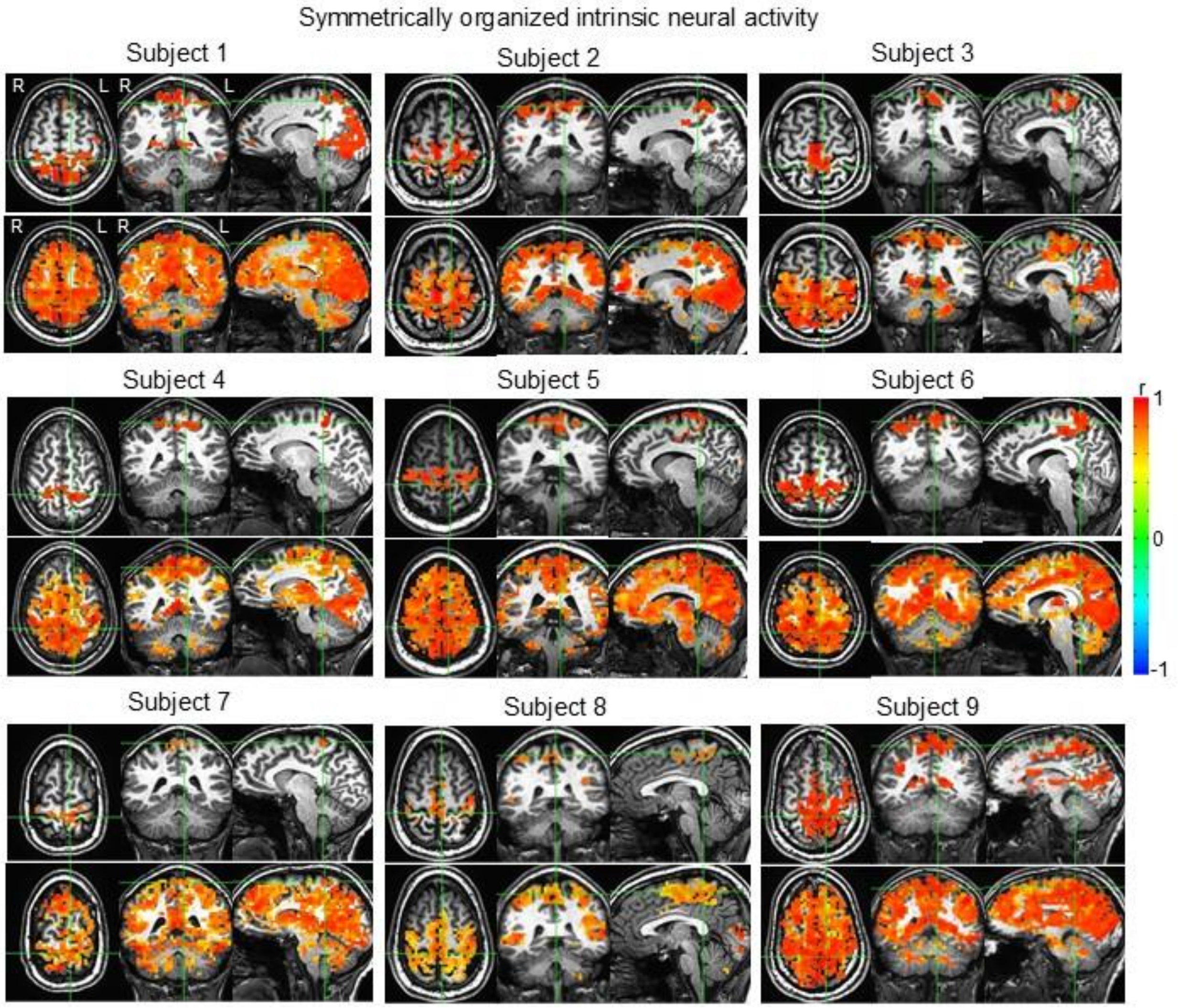

Suppl. Fig. 4 Illustration of the 1st identified symmetric FNN in the sensorimotor area (Fig. 5, top row) and its corresponding functionally connected FNNs for each subject. For each subject, the two green lines in each image of the top panel indicate the positions of the other two images, respectively. The crosspoint of these three lines indicates one of the two anatomic areas covered by that symmetric FNN. The functionally connected FNNs showed a substantial variation from subject to subject, demonstrating a high degree of individuality among these subjects.

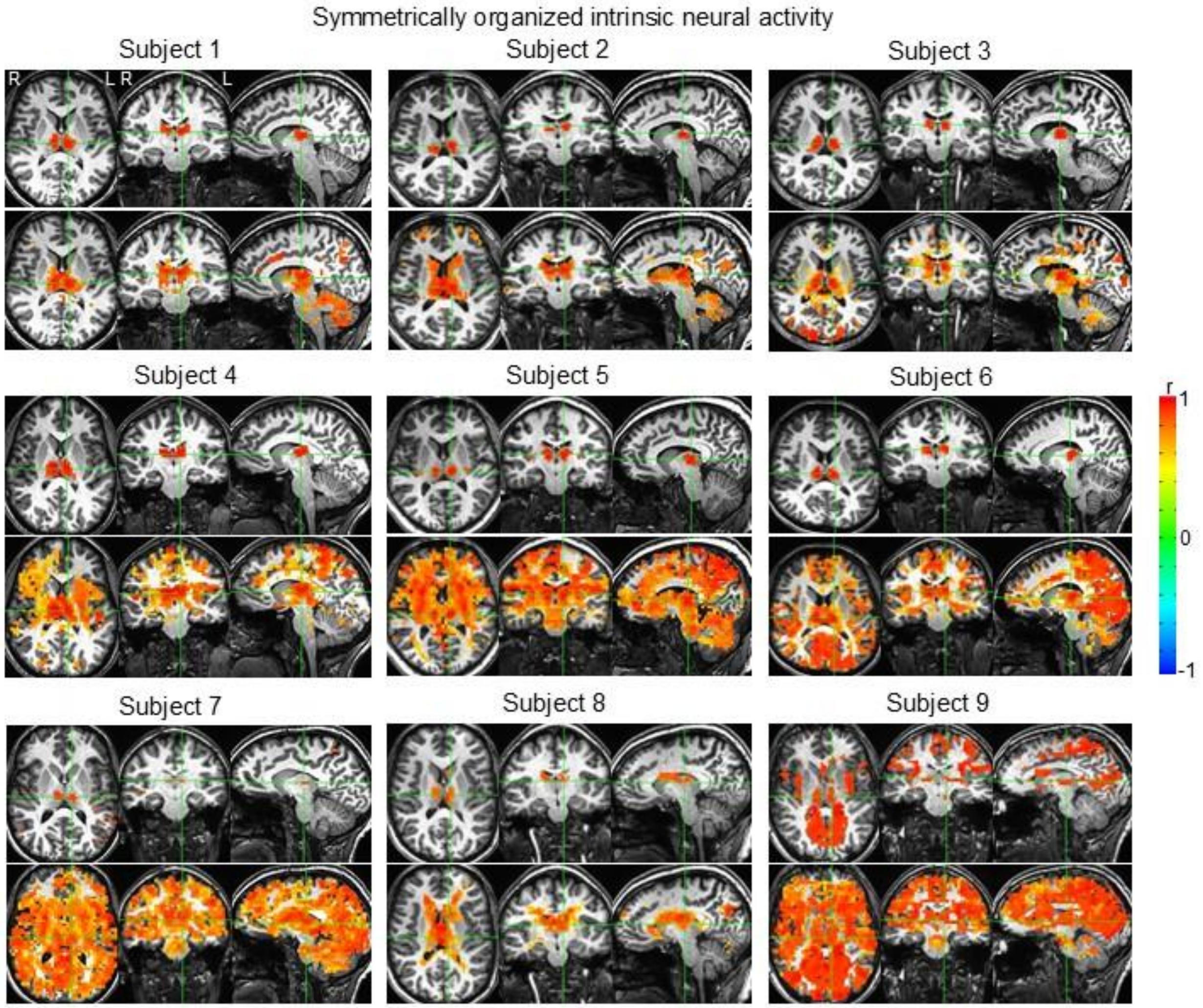

Suppl. Fig. 5 Illustration of the 2nd identified symmetric FNN in the thalami (Fig. 5, second row) and its corresponding functionally connected FNNs for each subject. For each subject, the two green lines in each image of the top panel indicate the positions of the other two images, respectively. The crosspoint of these three lines indicates one of the two anatomic areas covered by that symmetric FNN. Similarly to that in Suppl. Fig. 4, these functionally connected FNNs also showed a substantial variation from subject to subject, further demonstrating a high degree of individuality among these subjects.

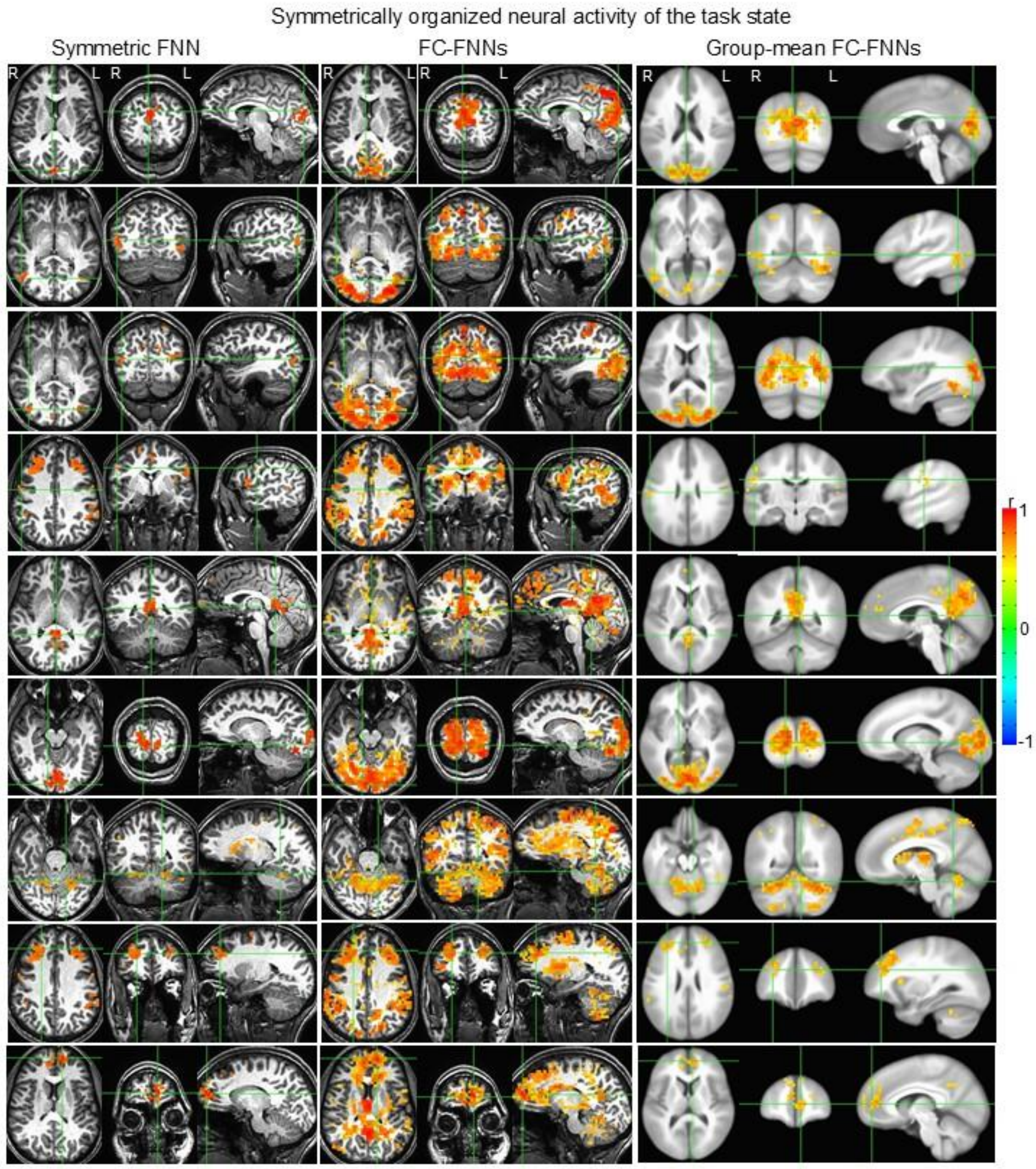

Suppl. Fig. 6 Illustration of the rest nine identified symmetric FNNs (left three columns) with their corresponding functionally connected (FC) FNNs (middle three columns) for the representative subject and group-mean FC FNNs averaged over the nine subjects (right three columns). For each grouped axial, coronal and sagittal images in the left three columns, the two green lines in each image indicate the positions of the other two images, respectively. The crosspoint of these three lines indicates one of the two anatomic areas covered by that symmetric FNN.

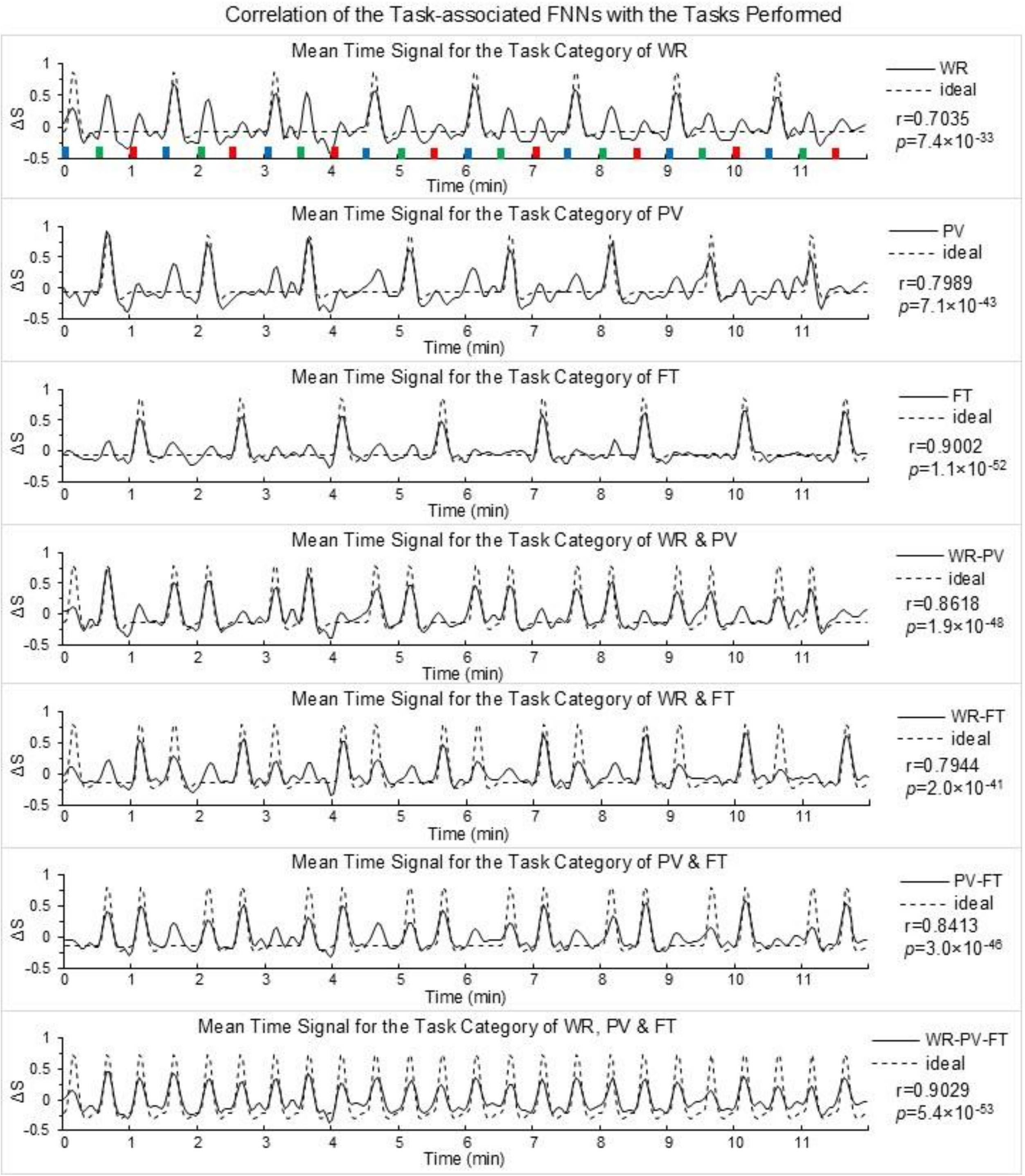

Suppl. Fig. 7 Validation of the identified task-associated FNNs with the tasks performed for each task category. The task paradigm consisted of 24 tasks shown by the 24 bars:

blue bars representing word-reading (WR), green bars pattern-viewing (PV) and red finger-tapping (FT). Each task lasted 6s followed by 24s rest, forming one task trial.
